# In Situ Electrochemical Studies of the Terrestrial Deep Subsurface Biosphere at the Sanford Underground Research Facility, South Dakota, USA

**DOI:** 10.1101/555474

**Authors:** Yamini Jangir, Amruta A. Karbelkar, Nicole M. Beedle, Laura A. Zinke, Greg Wanger, Cynthia M. Anderson, Brandi Kiel Reese, Jan P. Amend, Mohamed Y. El-Naggar

**Affiliations:** Department of Physics and Astronomy, University of Southern California, Los Angeles, California, USA; Department of Chemistry, University of Southern California, Los Angeles, California, USA; Department of Biological Sciences, University of Southern California, Los Angeles, California, USA; Department of Earth Science, University of Southern California, Los Angeles, California, USA; Center for the Conservation of Biological Resources, Black Hills State University, Spearfish, South Dakota, USA; Department of Life Sciences, Texas A&M University, Corpus Christi, Texas, USA

## Abstract

The terrestrial deep subsurface is host to significant and diverse microbial populations. However, these microbial populations remain poorly characterized, partially due to the inherent difficulty of sampling, *in situ* studies, and isolating of the *in situ* microbes. Motivated by the ability of microbes to gain energy from redox reactions at mineral interfaces, we here present *in situ* electrochemical colonization (ISEC) as a method to directly study microbial electron transfer activity and to enable the capture and isolation of electrochemically active microbes. We installed a potentiostatically controlled ISEC reactor containing four working electrodes 1500 m below the surface at the Sanford Underground Research Facility. The working electrodes were poised at different redox potentials, spanning anodic to cathodic, to mimic energy-yielding mineral reducing and oxidizing reactions predicted to occur at this site. We present a 16S rRNA analysis of the *in situ* electrode-associated microbial communities, revealing the dominance of novel bacterial lineages under cathodic conditions. We also demonstrate that the *in situ* electrodes can be further used for downstream electrochemical laboratory enrichment and isolation of novel strains. Using this workflow, we isolated *Bacillus*, *Anaerospora*, *Comamonas*, *Cupriavidus*, and *Azonexus* strains from the electrode-attached biomass. Finally, the extracellular electron transfer activity of the electrode-oxidizing *Comamonas* strain (isolated at −0.19 V vs. SHE and designated WE1-1D1) and the electrode-reducing *Bacillus* strain (isolated at +0.53 V vs. SHE and designated WE4-1A1-BC) were confirmed in electrochemical reactors. Our study highlights the utility of *in situ* electrodes and electrochemical enrichment workflows to shed light on microbial activity in the deep terrestrial subsurface.

**SIGNIFICANCE:** A large section of microbial life resides in the deep subsurface, but an organized effort to explore this deep biosphere has only recently begun. A detailed characterization of the resident microbes remains scientifically and technologically challenging due to difficulty in access, sampling, and emulating the complex interactions and energetic landscapes of subsurface communities with standard laboratory techniques. Here we describe an in situ approach that exploits the ability of many microbes to perform extracellular electron transfer to/from solid surfaces such as mineral interfaces in the terrestrial subsurface. By deploying and controlling the potential of in situ electrodes 4850 ft below the surface at the Sanford Underground Research Facility (South Dakota, USA), we highlight the promise of electrochemical techniques for studying active terrestrial subsurface microbial communities and enabling the isolation of electrochemically active microbes.

## INTRODUCTION

Minerals that contain redox active elements (e.g., S, Fe, Mn) are abundant in subsurface environments, and can support the growth of microorganisms by acting as electron acceptors for heterotrophic respiration or electron donors for lithotrophic, and often autotrophic, metabolism (1–7). This process of extracellular electron transfer (EET) to or from minerals is best characterized in a handful of Fe-reducing bacteria, especially *Geobacter* and *Shewanella* species (6, 8, 9). To overcome the hurdle of electron transfer across the otherwise electrically-insulating cell envelope, these bacteria rely on a number of mechanisms: multiheme cytochrome complexes that bridge the periplasm and outer membrane (10–12), microbial nanowires that reach out to distant electron acceptors (13, 14), and soluble redox shuttles that diffusively link cells to external surfaces (15, 16). EET can be electrochemically mimicked on electrode surfaces that function as surrogate electron acceptors (anodes) or donors (cathodes) to support microbial metabolism, depending on the poised potentials of these electrodes (17–22). As a result, electrochemical enrichments have been applied to microbial samples from a variety of environments (23–31). When combined with surveys of microbial community structure, these electrochemical techniques greatly expanded our understanding of the phylogenetic diversity of microbes capable of colonizing electrodes and led to the isolation of novel microorganisms capable of EET to anodes (32–37). While our mechanistic understanding of the molecular pathways that underlie inward EET (i.e., electron transfer from rather than to surfaces) lags behind metal reduction pathways, electrode-based techniques have highlighted the diversity of microbes capable of electron uptake from cathodes: acetogens, methanogens, as well as iron- and sulfur-oxidizers (38–45).

The Sanford Underground Research Facility (SURF), located in the Black Hills of South Dakota, USA is the site of the former Homestake Gold Mine. It is hosted in quartz-veined, sulfide-rich segments of an Early Proterozoic, carbonate-facies iron-formation (46–48). The facility’s tunnels provide access, maximally 4850 ft (1478 m) below ground surface (bgs), to deep subsurface fluids through a network of manifolds. The residence times of the accessible fluids range from very recent in the shallowest levels to much older (>10,000 years) fluids reaching the deeper levels (49, 50). The mine was converted to a state-run science facility in 2011, with emphasis on rare-process physics (51), and this accessible infrastructure provided a remarkable portal for *in situ* studies of the deep terrestrial biosphere. Osburn *et al*. (50) recently combined geochemical data, energetic modeling, and 16S rRNA gene sequences from different boreholes to both assess the *in situ* microbial communities and predict energy-yielding metabolisms for unknown physiotypes. Notably, abundant energy was predicted for microorganisms from a variety of reactions including sulfur oxidation, iron oxidation, and manganese reduction, which motivated us to apply electrode-based techniques to assess and enrich for microbes capable of EET.

Here we report the deployment and operation of an *in situ* electrochemical colonization (ISEC) reactor fed by a manifold at the 4850 ft (1478 m) level of SURF. The reactor contained four working electrodes, poised at potentials spanning reducing/cathodic to oxidizing/anodic, to act as an inexhaustible source or sink of electrons for capture of potential mineral oxidizing and reducing bacteria, with emphasis on mimicking sulfur oxidation, iron oxidation, and manganese reduction predicted as energy-yielding reactions in this manifold (50). Our goals were twofold: 1) to demonstrate the use of electrodes as *in situ* observatories for directly detecting and characterizing microbial electron transfer activity, and 2) to capture important bacterial lineages active in the deep subsurface. Further, we utilized these *in situ* electrodes for subsequent laboratory enrichment and isolation of novel strains, and directly tested the electrochemical activity of a cathode-oxidizing *Comamonas* isolate and an anode-reducing *Bacillus* isolate. To our knowledge, this effort represents the first potentiostatically-controlled multi-electrode reactor deployed *in situ* in a deep subsurface environment.

## MATERIAL AND METHODS

### Field Measurements

The effluent was sampled from an exploratory borehole (DUSEL 3A) located at the 1478 m (4850 ft) level close to the Yates shaft at SURF in Lead, South Dakota (USA). The 214 m-long borehole, drilled in 2009 intersects the Precambrian amphibolite metamorphic schists (46, 47) and has been capped with manifolds to prevent flooding. Initial aqueous chemistry of DUSEL 3A fluid was obtained for thermodynamic modeling in February 2014 (50). Further, detailed geochemical data were acquired during installation (December 2014) and conclusion (May 2015) of the ISEC reactor incubation. Aqueous chemistry (Table 1) was measured as described earlier (50). Briefly, oxidation-reduction potential (ORP), conductivity, pH, temperature, and total dissolved solids (TDS) were measured *in situ* with an Ultrameter II 6PFCE (Myron L Company, Carlsbad, CA). The redox-sensitive species were measured using Hach DR/2400 portable field spectrophotometers and associated reaction kits (Hach Company, Loveland, CO). Major anions and cations were measured using ion chromatography with Metrohm column Metrosep A SUPP 150 and Metrosep C6 - 250/4.0, respectively (Metrohm, Fountain Valley, CA).

**Table 1:** Chemical composition of fluid from DUSEL 3A at the 4850 ft level, Sanford Underground Research Facility (SD, USA), after commissioning and decommissioning of in-situ electrochemical colonization (ISEC) reactor.

|  | **Dec 2014** | **May 2015** |
| --- | --- | --- |
| pH | 7.08 | 7.75 |
| Temperature (°C) | 18.7 | 19.31 |
| Conductance (μS/cm) | 6514 | 7869 |
| TDS (ppm) | 3257 | 3929 |
| ORP (mV) | 59.2 | -50 |
| **Dissolved Constituents** |  |  |
| dO<sub>2</sub> (mg/L) | 3.79 | 0 |
| S<sup>2-</sup> (mg/L) | 17 | 17 |
| Fe<sup>2+</sup>(mg/L) | 0.6 | 0.6 |
| NO<sub>3</sub><sup>-</sup>(mg/L) | 0.4 | 0.2 |
| NH<sub>3</sub> (mg/L) | 0.11 | 0.07 |
| SiO<sub>2</sub> (mg/L) | 24.7 | 9.7 |
| Mn<sup>2+</sup> (mg/L) | 0.4 | 0.9 |
| PO<sub>4</sub><sup>3-</sup> (mg/L) | 1.24 | 1.09 |
| NO<sub>2</sub><sup>-</sup> (mg/L) | BDL | 0.001 |
| Br<sub>2</sub> (mg/L) | BDL | 0.01 |
| Li<sup>+</sup> (mg/L) | 0.024 | 0.035 |
| Na<sup>+</sup> (mg/L) | 10.619 | 5.698 |
| K<sup>+</sup> (mg/L) | 0.229 | 0.751 |
| Cl<sup>-</sup> (mg/L) | 0.316 | 0.385 |
| SO<sub>4</sub><sup>2-</sup> (mg/L) | 14.049 | 46.696 |

### In Situ Electrochemical Colonization Reactor

Fluid from DUSEL 3A was pumped through the ISEC reactor (Figure 1) for 5 months via peristaltic pumps (MasterFlex L/S Digital Drive, EW-07522-20, Cole-Parmer, USA) at a flow rate of 1 mL/min, providing a dilution rate of 1 day-1. Aqueous chemistry (ORP, conductivity, pH, and temperature) of the ISEC effluent was logged using a multiparameter HI9829 meter (Hanna Instruments, Woonsocket, RI). ISEC is a standard electrochemical half-cell design, custom fabricated from a 1 L spinner flask (CLS-1410, Chemglass Life Sciences, Vineland, NJ) to incorporate an Ag/AgCl reference electrode (3M NaCl, MF-2052, BASi, West Lafayette, IN) and a platinum counter electrode (CHI115, CH Instruments, Austin, TX). It holds a PTFE assembly of five threaded rods, each of which supported a working electrode (WE) composed of 6 cm × 7 cm carbon cloth (PW06, Zoltek, St. Louis, MO, USA). During enrichment, four working electrodes (WE1, WE2, WE3, and WE4) were poised at −0.19, +0.01, +0.26 and +0.53 V vs. SHE, respectively, using a four-channel potentiostat (EA164 Quadstat, EDaq, USA). Potentials were chosen to mimic conditions consistent with elemental sulfur oxidation (−0.19 V vs. SHE), iron oxidation (+0.01 V vs. SHE), and manganese reduction (+0.53 V vs. SHE), all predicted as putative energy-yielding metabolisms for the microbial community inhabiting the DUSEL 3A fluid (50). Since the reduction potential of many cytochromes ranges from −0.5 to +0.3 V vs. SHE (52) a fourth potential was also applied at +0.26 V vs. SHE. All electrical connections were made using titanium insulated wire (KeegoTech, Menlo Park, CA, USA). The complete set-up was autoclaved along with the working and the counter electrodes. The reference electrode was ethanol-sterilized before it was inserted. Applied redox potentials and current production were controlled and recorded via the eCorder eCHART software (eDAQ Inc., Colorado Springs, CO). Before the deployment of the ISEC reactor in December 2014, the SURF facility made modifications to the manifold’s design, which could potentially lead to backflow of fluid from a dehumidifier into the reactor. To avert this contamination, in December 2014, a check valve was installed upstream of the solenoid valve as shown in the schematic diagram of ISEC reactor (Fig 1). The reactor was incubated with a continuous flow of DUSEL 3A fluid for 5 months beginning December 2014, monitored monthly for proper operation, and finally retrieved in May 2015. During decommission, small sections of carbon cloth from each potential were collected for microscopy, active microbial community analysis, and further laboratory electrochemical enrichments as described below.

**Figure 1:**
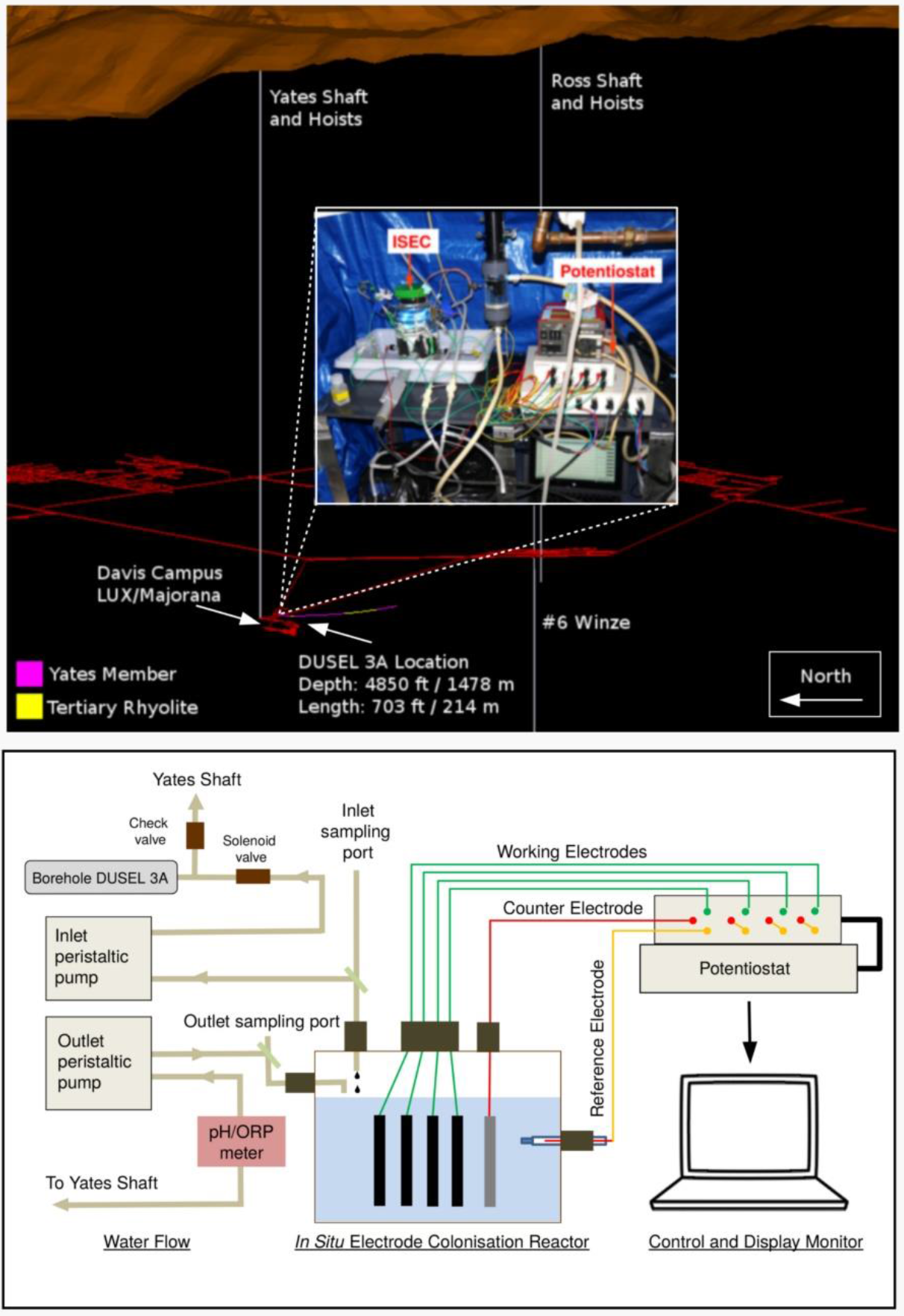
(Top) A 3-dimensional layout of Sanford Underground Research Facility (image to scale). DUSEL 3A, located at 4850 ft level close to Davis Campus cutting the iron-rich Precambrian Yates member, was selected as the site for *in situ* electrochemical colonization (ISEC) reactor deployment. An image of the ISEC reactor after deployment (Dec 2014) is shown in the inset. (Bottom) Schematic diagram of ISEC reactor. The four carbon cloth working electrodes were poised at −0.19, +0.01, +0.26 and +0.53 V vs. SHE, respectively, using a 4-channel potentiostat to enrich for electrode oxidizing and reducing microbes in a single reactor. DUSEL 3A fluid was fed directly into the reactor and the effluent was pumped out to Yates shaft.

### Secondary Laboratory Electrochemical Enrichment

At the conclusion of ISEC deployment, the carbon cloth working electrodes (3 cm × 6 cm) were placed into autoclaved 125 mL serum bottles containing filtered (0.22 μm) DUSEL 3A fluid filled to capacity and sealed with sterile butyl rubber stoppers and aluminum seals (Chemglass Life Sciences, USA). The serum bottles were stored in vacuum-sealed Mylar bags (ShieldPro, USA) with oxygen absorbers (OxyFree, USA) to minimize oxygen exposure during transportation. In the laboratory, secondary electrochemical enrichments were performed using the poised carbon cloth from the reactor in four separate three-electrode electrochemical cells (50 mL total volume). A standard three-electrode glass cell comprised of: 1) working electrode (WE): a carbon cloth with dimensions 1 cm × 1 cm; 2) counter electrode (CE): a platinum wire (CHI115, CH Instruments, USA), and 3) a reference electrode (RE): 1 M KCl Ag/AgCl reference electrode (CHI111, CH Instruments, USA). Specialty media was designed to mimic the *in situ* aqueous chemistry of DUSEL 3A at SURF, which included basal salts (L−1): NaCl (0.029 g), KH_2_PO_4_ (0.041 g), NH_4_Cl (0.160 g), FeSO_4_.7H_2_O (0.042 g), Na_2_SO_4_ (0.014 g), and amended with yeast extract (0.5 g), vitamins, and trace minerals (53). Media was maintained at pH ~7.5 using phosphate buffer (10 mM). Enrichment of electrode oxidizers was performed by augmenting SURF base media with sodium bicarbonate (10 mM), poising electrodes at −0.19 and +0.01 V vs. SHE, and continuous purging of a gas mix (CO_2_: air v:v 20:80) in the electrochemical cell. To enrich for electrode reducers, media was augmented with sodium acetate (5 mM), electrodes were poised at +0.26 and +0.53 V vs. SHE, and continuous N_2_ gas purging was used to maintain anoxic conditions. These secondary electrochemical enrichments were performed in batch mode (media changed every 7-10 days) for 2 months.

### Active Microbial Community Analysis

The active microbial community was analyzed by extracting 16S ribosomal RNA (rRNA) transcripts from: 1) dehumidifier fluid (as a potential contaminant) (2 L), 2) DUSEL 3A fluid (2 L), 3) electrode-attached biofilms in ISEC (3 x 4 cm), 4) electrode-attached biofilms of the laboratory electrochemical enrichments (1 x 0.5 cm), and 5) a blank sample containing no sample. The DUSEL 3A and dehumidifier fluid were filtered through 0.22 μm sterivex filters (Millipore, USA) to collect microbiological samples. All microbiological samples were immediately placed on dry ice for transportation to the University of Southern California and kept in a −80 °C freezer in the laboratory until extraction. Total RNA was extracted from the filters and electrode-attached biofilms (ISEC and secondary laboratory electrochemical enrichments) via physical (freeze, thaw, vortex), chemical (sodium dodecyl sulfate and EDTA), and biological (lysozyme) disruption of the cell wall prior to phenol-chloroform extraction as described previously (54). A control that included only reagents and no sample material was extracted alongside the samples to confirm lack of contamination. Any residual DNA was digested using TURBO DNA-free TM Kit (ThermoFisher Scientific, USA) and verified to be free of DNA by running a PCR of the RNA extraction. Samples were reverse transcribed with MMLV (Promega, USA) using 806R primer (55). The complementary DNA (cDNA) of each sample was sent to Molecular Research DNA (Shallowater, TX, USA) for library preparation and high throughput sequencing. Sequencing library preparation included amplifying the hypervariable region V4 of the bacterial 16S rRNA transcript using barcoded fusion primers 515F and 806R (55), according to previously described methods (Dowd et al., 2008). Following amplification, all PCR products from different samples were mixed in equal concentrations and purified using Agencourt Ampure beads (Agencourt Bioscience Corporation, MA, USA). Samples were sequenced on Roche 454 FLX titanium platform using recommended reagents and following manufacturer’s guidelines.

The sequences were processed using mothur pipeline (56). The barcodes and the primers were removed from each sequence, followed by trimming to remove any ambiguous base calls, average quality scores less than 20, and homopolymer runs more than 8 bp. A total of 422,490 high-quality unique sequences remained and the resulting average sequence length was 260 bp for all samples. The trimmed sequences were aligned with the SILVA-based reference alignment (57) using the Needleman-Wunsch pairwise alignment method (58). Chimeras were removed using UCHIME (59), and a distance matrix was created. The sequences were clustered to identify unique operational taxonomical units (OTUs) at the 97% level, followed by creating a shared OTU file amongst various samples, and the taxonomy was assigned using mothur formatted Ribosomal Database Project (RDP) training set version 16. The resultant microbiome was processed using “Phyloseq”, “Microbiome” and “Vegan” package available in R (version 3.2.2). The dataset (OTU shared file, taxonomy file, and the metadata file) were introduced as a phyloseq objects in R. The microbiome data was pre-processed by removing contaminant OTUs from the extraction reagent. OTUs not represented in the DUSEL 3A (source fluid) were also removed from the dataset to account for any contamination from initial dehumidifier backflow. The resulting filtered microbiome data (including absolute abundance data from DUSEL 3A, ISEC, and Laboratory enrichments) was used to calculate the diversity indices, MDS plots, and statistics including Adonis and betadisper. The absolute abundance microbiome data of m phylotype/OTUs and n samples (*C* = (*c*_*ij*_) ∈ ℕ^*m*×*n*^, where *c*_*ij*_ are the number of reads for a phylotype/OTU (i) in the sample (j)) was transformed into the relative abundances by total sum scaling 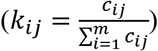. Next, to study the variation of relative abundance of a phylotype/OUT across sample the dataset was normalized to the maximum 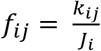, *where J_i_*=*maximum*{*k*_*ij*_}. The raw sequencing data have been uploaded to National Center for Biotechnology Information (NCBI) Sequence Read Archive (SRA) database (accession number: SRR8537881-SRR8537882) under BioProject PRJNA262938.

### Isolation and Electrochemical Measurements of Pure Isolates

Electrode-attached biomass from each of WE1 and WE4 was streaked on R2A agar plates (60) and incubated at 30°C to obtain single colonies. Morphologically distinct colonies were re-streaked on fresh R2A agar plates, resulting in multiple isolates. For taxonomic classification, isolates were grown to late exponential phase in liquid R2A at 30°C and DNA extraction performed using the UltraClean^®^ Microbial DNA Isolation kit (Mo Bio Laboratories, Carlsbad, CA, USA). The bacterial 16S rRNA gene was PCR-amplified from the extracted DNA using primers 8F and 1492R (Integrated DNA Technologies, USA). The purified PCR product (PureLink PCR Purification Kit, Life Technologies, CA, USA) was sent for sanger sequencing to Genewiz (South Plainfield, NJ, USA) from the 1492R primer. The 600 bp length sequences of the isolated strains have been deposited to NCBI SRA (accession numbers: MK483257, MK483258). The chronoamperometry (i.e., current vs. time at a fixed potential) measurement of each isolate was performed in a standard three-electrode glass electrochemical cell (50 mL volume) in triplicates.

*Comamonas* sp. (designated strain WE1-1D1) was grown aerobically to stationary phase from frozen stocks (−80 °C in 20% glycerol) in R2A media and the culture was used to inoculate 750 mL of SURF base medium at 1% (v:v). After reaching the mid-exponential phase, the culture was pelleted by centrifugation at 5369 × g for 10 min, washed three times and re-suspended in 15 mL fresh SURF base medium without electron donor or trace minerals. 5 mL of this suspension was introduced into three separate electrochemical cells, already containing 50 mL SURF base medium, after the abiotic current was stabilized to a steady baseline. The electrochemical cell was continuously purged with 100% purified compressed air, and chronoamperometry was performed with the working electrode poised at −0.012 V vs. SHE. After cyclic voltammetry was completed at a scan rate of 1 mV/s, chronoamperometric measurement was resumed at −0.012 V vs. SHE, and ~10.7 mM potassium cyanide was added after ~1.75 hours. Cell densities were determined by plate counts.

*Bacillus* sp. (designated strain WE4-1A1-BC) was grown aerobically to stationary phase from frozen stocks (−80 °C in 20% glycerol) in R2A media and the culture was used to inoculate 500 mL of SURF base medium at 1% (v:v). After reaching mid-exponential phase, the culture was pelleted by centrifugation at 7000 × g for 10 min, washed two times and resuspended in 10 mL fresh SURF base medium without electron donor. A total of 5 mL of the suspension was introduced into the electrochemical cell, containing 50 mL SURF base medium augmented with sodium acetate (5 mM), after the abiotic current was stabilized to a steady baseline. The electrochemical cell was continuously purged with N_2_ to maintain anaerobic conditions, and chronoamperometry was performed with the working electrode poised at +0.53 V vs. SHE. Cell densities were determined by plate counts. The reproducibility of the experiment was tested in triplicates.

### Microscopy

For scanning electron microscopy (SEM), electrode samples were fixed overnight in 2.5% glutaraldehyde. Samples were then subjected to an ethanol dehydration series (25%, 50%, 75%, 90%, and 100% v/v ethanol for 15 min each treatment) and critical point drying (Autosamdri 815 critical point drier, Tousimis Inc., Rockville, MD, USA). The samples were then mounted on aluminum stubs, coated with gold (Sputter coater 108, Cressington Scientific), and imaged at 5 keV using a JEOL JSM 7001F low vacuum field emission SEM.

## RESULTS AND DISCUSSION

### *In situ* current response and active microbial community composition

The ISEC working electrode potentials were chosen to mimic conditions similar to elemental sulfur oxidation (WE1 = −0.19 V vs. SHE), iron (II) oxidation (WE2 = +0.01 V vs. SHE), and manganese (IV) reduction (WE4 = +0.53 V vs. SHE), all predicted as putative energy-yielding metabolisms for the microbial community inhabiting DUSEL 3A fluid (50). Since the reduction potential of many cytochromes ranges from −0.5 to +0.3 V vs. SHE (52), a fourth potential was also applied at WE3 = +0.26 V vs. SHE.

The reduction potentials were evaluated by: 1) converting the standard Gibbs free energy (∆G^0^) to standard reduction potential 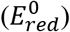 of the reaction via 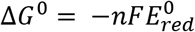 (where: n refers to the number of electron transferred and F stands for the Faraday constant), and 2) using the geochemical data of DUSEL 3A liquid to evaluate *in situ* reduction potential (*E*_*red*_) via the Nernst Equation 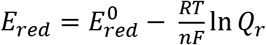 (where R refers to the gas constant, T denotes the temperature in kelvin, n refers to the number of electron transferred and F stands for the Faraday constant and *Q*_*r*_ represents the reaction quotient of the cell reaction). Large negative currents were detected on both WE1 and WE2, −276.68 ± 43.80 μA and −168.76 ± 26.82 μA respectively, consistent with the expected cathode oxidation conditions. While significantly smaller in magnitude, cathodic currents were also detected on WE3 and WE4 at higher (nominally anodic) potentials (Table 2). However, it is important to note that the currents observed in the ISEC reactor are a combination of both biotic and abiotic redox reactions occurring on electrode surfaces. Oxygen leakage into the reactor and associated oxygen reduction currents can, therefore, play a role. In May 2015 (during decommissioning of the ISEC reactor), ORP values of DUSEL 3A fluid and the effluent of the ISEC reactor were −50 mV (Table 1) and 200 mV (Table 2), respectively; indicating oxygen leakage inside the ISEC reactor.

**Table 2:**
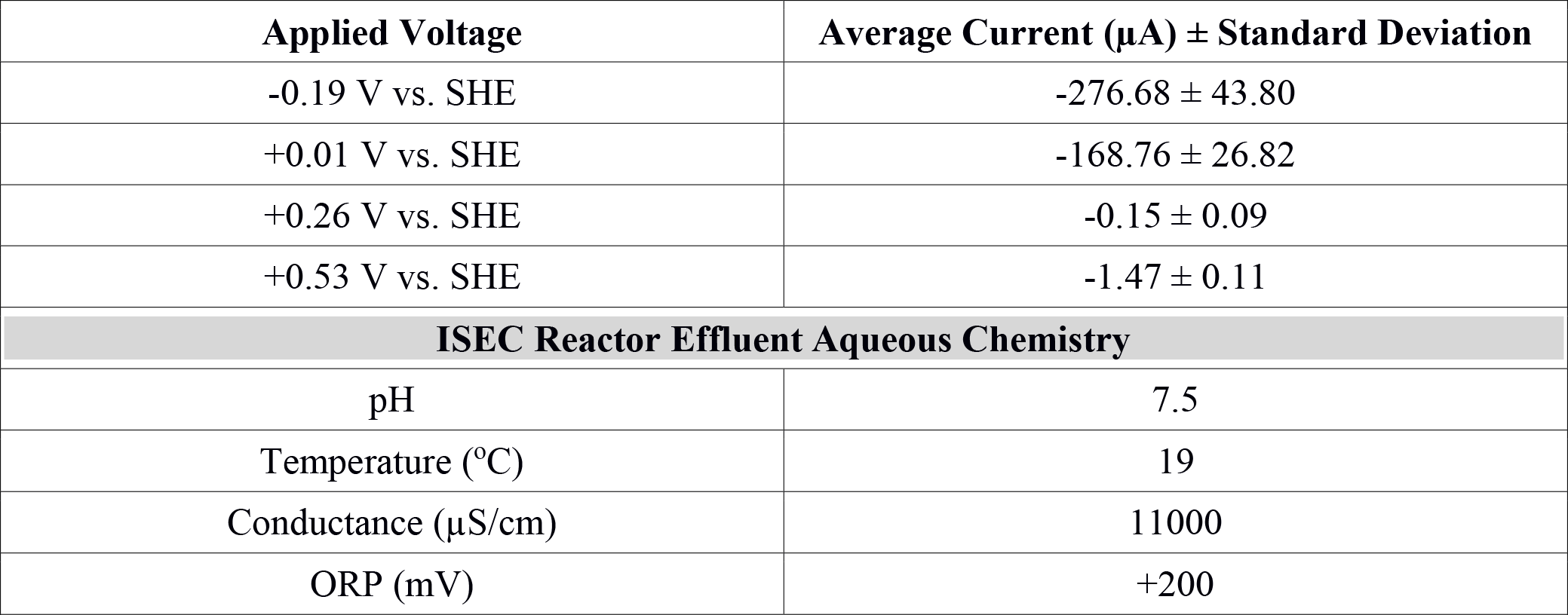
Mean current (last 40 days) observed at various redox potentials in the *in situ* electrochemical colonization (ISEC) reactor and aqueous water chemistry of ISEC reactor effluent.

| Applied Voltage | Average Current ($\mu\text{A}$) ± Standard Deviation |
| --- | --- |
| -0.19 V vs. SHE | $-276.68 \pm 43.80$ |
| +0.01 V vs. SHE | $-168.76 \pm 26.82$ |
| +0.26 V vs. SHE | $-0.15 \pm 0.09$ |
| +0.53 V vs. SHE | $-1.47 \pm 0.11$ |
| **ISEC Reactor Effluent Aqueous Chemistry** |  |
| pH | 7.5 |
| Temperature ($^{\circ}\text{C}$) | 19 |
| Conductance ($\mu\text{S}/\text{cm}$) | 11000 |
| ORP (mV) | +200 |

Nucleic-acid analysis has proven to be effective for characterizing the phylogenetic, taxonomic and functional structure of microbial assemblages. Since extracted DNA may originate from extracellular DNA pools, dead, and/or dormant cells, sequenced 16S rRNA genes represent the total microbial community rather than its active fraction. On the other hand, ribosomal RNA transcripts describe the active portion of the microbial community (61) and hence, an RNA-based molecular approach was chosen for this study. Herein we present the taxonomic assignments in terms of relative percent of classified sequences with respect to the total number of sequences.

The *in situ* microbial community analyses determined by pyrosequencing the 16S rRNA transcripts, revealed that Bacteria dominated DUSEL 3A fluid (99.65%). The bacterial sequences most closely matched the phyla *Proteobacteria* (83.23%), *Bacteroidetes* (10.14%), *Firmicutes* (0.50%), *Chloroflexi* (0.13%), *Planctomycetes* (0.003%). Some sequences (5.82%) did not match to any phyla and have been labeled as unclassified. Within *Proteobacteria*, the majority of sequences were identified as *Gammaproteobacteria* (26.03%), followed by unclassified *Proteobacteria* class (24.45%), *Deltaproteobacteria* (17.78%), *Alphaproteobacteria* (10.30%) and *Betaproteobacteria* (4.66%) (Figure 2). The archaeal community in DUSEL 3A fluid was dominated by sequences related to *Crenarchaeota* (0.19%), *Euryarchaeota* (0.13%), and *Thaumarchaeota* (0.03%) (Figure 2). In a DNA-based study, Osburn et al. (2014) also observed a predominance of Bacteria over Archaea in the *in situ* fluid at the 4850-ft level of SURF. They further observed a higher abundance of *Firmicutes*, which we attribute to the DNA-versus RNA-based microbial community analysis because this phylum has been shown to experience dormancy through sporulation.

**Figure 2:**
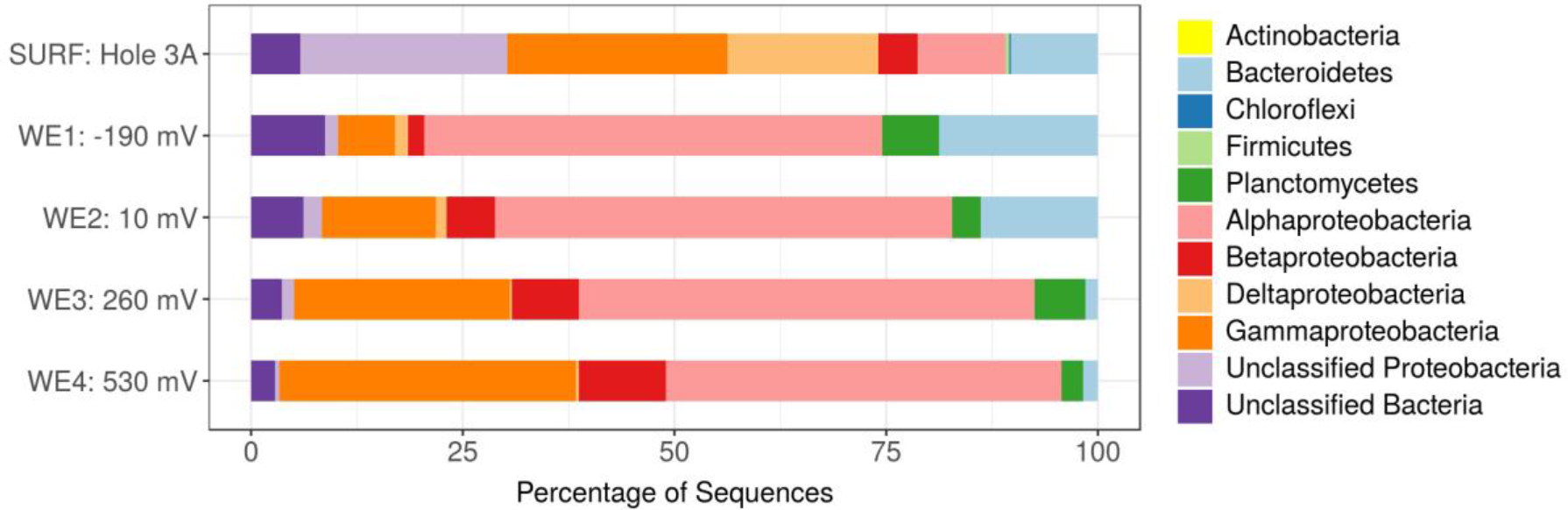
Phylum classification of the active *in situ* bacterial community of DUSEL 3A fluid (labeled SURF: Hole 3A) and the four separate working electrodes in the *in situ* electrochemical colonization (ISEC) reactor (labeled WE1-WE4). The unclassified (e.g., putatively novel microbiota) were relatively more abundant at the reducing potential (−0.19 V vs. SHE) where bacteria use the electrode as the electron donor.

Importantly, the active microbial community structure revealed some shifts between the DUSEL 3A *in situ* fluid and the electrode-attached biofilms of the ISEC reactor. The majority of sequences obtained from electrode-attached biomass of ISEC working electrodes were most closely related to the *Proteobacteria*, specifically the A*lphaproteobacteria* (46.75-54.03%), *Gammaproteobacteria* (6.68-35.96%), *Betaproteobacteria* (1.98-10.24%), unclassified (0.57-2.17%), and *Deltaproteobacteria* (0.29-1.54%). The increased relative abundance of sequences identified as *Gammaproteobacteria* and *Betaproteobacteria* was observed with increasing electrode potential. The overall relative abundance of sequences related to *Planctomycetes* phylum substantially increased from the DUSEL 3A fluid (0.003%) to ISEC working electrodes (3-6%). On the other hand, the relative abundance of sequences identified as *Chloroflexi* and *Firmicutes* phyla decreased from 0.13% and 0.49% to 0.001% and 0.002% respectively. The relative abundance of sequences related to *Bacteroidetes* from the electrode-attached biofilm (across different working electrodes) of the ISEC reactor ranged from 1.45-18.76% and unclassified OTUs at the phyla level ranged from 2.83-8.72%. The ISEC reactor captured the majority of the bacterial community, but some members, including *Methanobacteria*, unclassified *Clostridia*, *Methylophilaceae*, and *Moraxellaceae*, were not represented. These populations were originally present in low abundances (less than 0.13%) in DUSEL 3A fluid. Suspected oxygen leakage into the ISEC reactor potentially hindered the survival of obligate anaerobes belonging to the class *Methanobacteria* and *Clostridia* (62). Many bacterial lineages belonging to unclassified (at the phylum level), *Bacteroidetes* and *Alphaproteobacteria* favored the lower potential electrodes; this observation highlights the potential for largely uncharacterized microbial lineages to perform extracellular electron uptake in subsurface environments. Within the sequences classified as Archaea, the phylum *Thaumarchaeota* (ammonia-oxidizing *Candidatus Nitrosoarchaeum*) was identified on all ISEC working electrodes. However, *Crenarchaeota* sequences (0.002%) were only present at an electrode potential of −0.19 V.

Based on the 16S rRNA sequencing data, the bacterial communities of the electrode attached biofilms demonstrated distinct potential-dependent distribution patterns (Figure 3). Below we discuss these patterns in light of previously reported physiotypes. However, we stress that the presence of specific lineages on electrodes is not in itself evidence of EET, since poised electrodes may support microbial consortia capable of diverse metabolisms, including syntrophy, heterotrophy or fermentation, depending on the microenvironment formed at the electrodes. This makes isolation and electrochemical characterization fundamental to confirming and investigating EET processes from the electrode-attached biofilm community.

**Figure 3:**
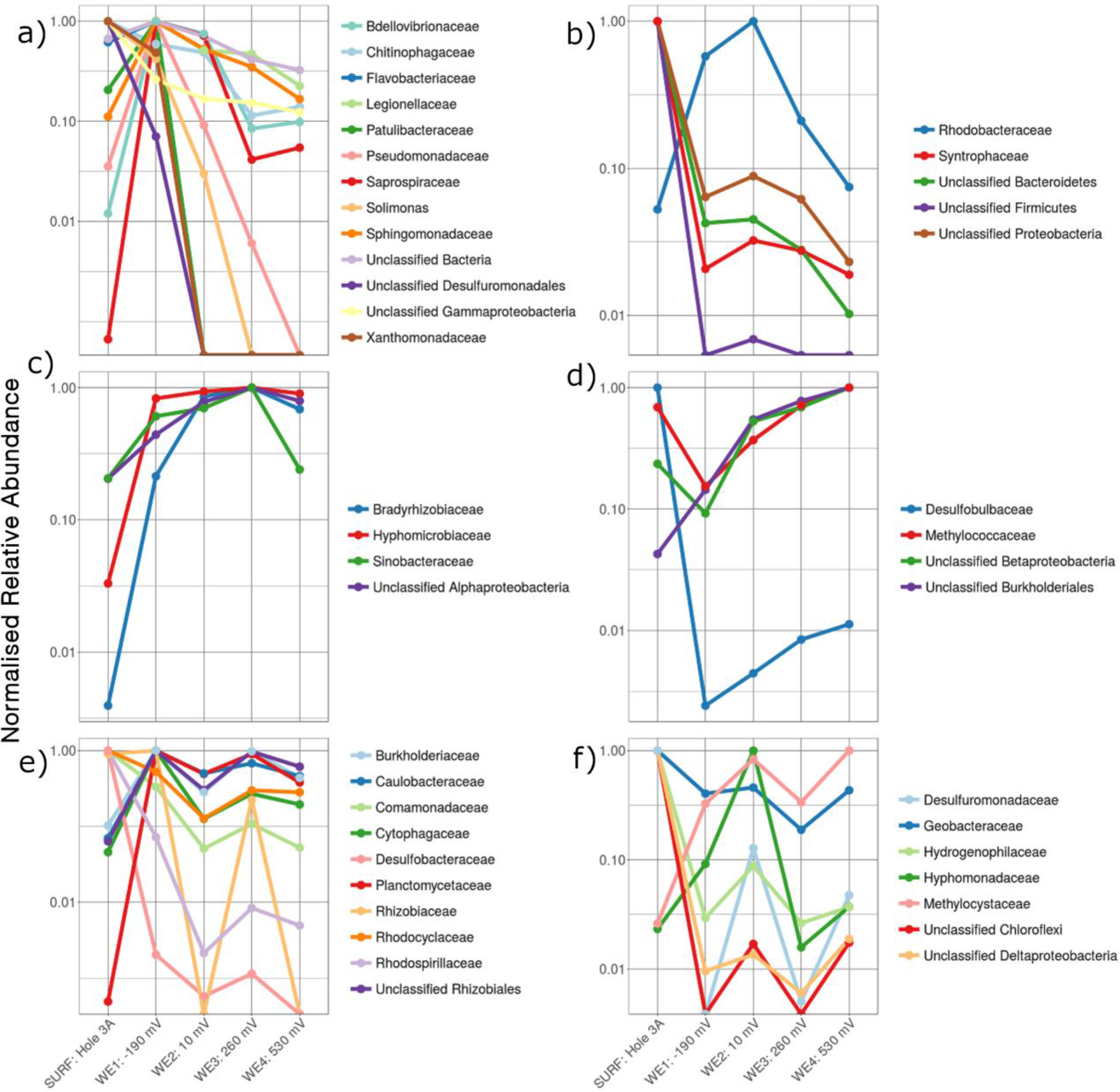
Differential normalized relative abundance of family-level active bacterial community sorted by poised electrodes of the *in situ* electrochemical colonization (ISEC) reactor in reference to the inoculum – fluid of SURF DUSEL 3A (labeled as SURF: Hole 3A).

At a potential of −0.19 V (WE1), chosen to mimic sulfur oxidation, the electrode was dominated by sequences closely related to the families *Bdellovibrionaceae, Chitinophagaceae, Flavobacteraceae, Legionellaceae, Patulibacteraceae, Psuedomonadaceae, Saprospiraceae, Solimonas, Sphingomonadaceae, Xanthomonadaceae*, and unclassified *Desulfuromonadales* (Figure 3a). Consistent with this, members of *Flavobacterium* (within the *Flavobacteraceae* family) have been shown to attach to and metabolize sulfur minerals, and this has been postulated to create micro-oxic environments for sulfur oxidizers (63). Some *Flavobacteria* genera have also co-existed with mat-forming neutrophilic iron oxidizers (64), and some Mn-oxidising strains belonging to *Flavobacterium* have been isolated from caves of upper Tennessee river basin (65). The family *Pseudomonadaceae* contains the metabolically diverse *Pseudomonads*, which has been studied inhabit varied niches. Marine and freshwater *Pseudomonads* have been shown to be capable of growing by oxidizing thiosulfate to tetrathionate (66). A *Pseudomonas/Pseudoalteromonas*-like *Gammaproteobacteria* was previously shown to putatively catalyze iron(II) oxidation under micro-aerobic conditions, isolated from volcanic seamount (67), which may contribute to the formation of iron mats in the deep marine subsurface. Other lineages captured at this potential included *Patulibacteraceae, Saprospiraceae, Chitinophagaceae, Sphingomonadaceae*, and *Solimonas*, each putatively containing aerobic respiratory metabolism among their members (68–73).

The potential +0.01 V vs. SHE (WE2), chosen to mimic iron oxidation, favored the colonization of members from bacterial families *Rhodobacteracea*e, *Syntrophaceae*, and many OTUs were identified as unclassified at the phyla level (Figure 3b). Some members of the family *Rhodobacteraceae*, such as gram-negative *Rhodobacter ferrooxidans* strain SW2, have been isolated from freshwater and characterized as photoautotrophic and Fe(II)-oxidizing (74). The *Rhodobacteraceae* family has been widely studied to understand banded iron formation in early Earth (75, 76). The abundance of many unclassified from the *Desulfuromonadales*, *Gammaproteobacteria*, *Bacteroidetes*, *Firmicutes*, and *Proteobacteria* at the lower electrode potentials highlights the importance of alternate sources of reducing power available in the subsurface. The families *Bradyrhizobiaceae* and *Hyphomicrobiaceae*, were predominantly abundant on WE3 poised at +0.26 V vs. SHE (Figure 3c). In previous studies, *Rhodopseudomonas palustris* strain TIE-1 (77), a phototrophic iron-oxidizing bacteria at circumneutral pH, was demonstrated to uptake electron from electrodes poised at +0.1 V vs. SHE (39). Sequences related to mesophilic sulfate-reducing *Desulfobulbaceae* (78) and type 1 methanotrophic *Methylococcaceae* (X.X%) (79) were relatively dominant on WE4 poised at +0.53 V vs. SHE, which was chosen to mimic manganese reduction (Figure 3d). Previous studies have also shown that members of the *Desulfobulbaceae* family, specifically *Desulfobulbus propionicus*, may putatively oxidize inorganic sulfur compounds coupled with the reduction of Mn(IV) (80). *D. propionicus* can also oxidize elemental S0 with an electrode (poised at +0.522 V vs. SHE) serving as the electron acceptor (81).

Some microbes adapt to broad ranges of environments (“habitat generalists”) as their key survival strategy, while others specialize in certain niches (“habitat specialists”) (82). In this study, we observed that some bacterial populations colonized working electrodes specific to a particular habitat - low and high electrode potential (Figure 3e and 3f, respectively). Among these, members of the bacterial families *Comamonadaceae*, *Rhodocyclaceae*, *Burkholderiaceae*, *Caulobacteraceae, Hyphomonadaceae, Hydrogenophilaceae*, *Desulfobacteraceae*, *Desulfuromonadaceae*, *Geobacteraceae*, have been shown to interact with minerals (83). *Rhodoferax ferrireducens* sp. Strain T118^T^, a member of the *Comamonadaceae* family, can support growth by oxidizing acetate with iron(III) reduction (84). Also, novel strains from the *Delftia* genus (*Comamonadaceae*) and *Azonexus* (*Rhodocyclaceae*) were demonstrated to donate electrons to poised electrodes (+0.522 V vs. SHE) in anaerobic conditions (34). A member of the *Burkholderiaceae* family, metallophilic gram-negative *Cupriavidus metallidurans*, has demonstrated to actively biomineralized particulate Au (85). *Desulfuromonadaceae* family contains members are capable of sulfur and Fe(III) reduction (86), and more recently *Desulfuromonas soudanensis* WTL was isolated using electrochemical enrichments from anoxic deep subsurface brine (32) with 38 multiheme cytochrome. In this study, the relative predominance of *Desulfuromonadaceae* at lower and higher potentials indicates their ability to survive/grow in niches (Figure 3f). Finally, *Geobacteraceae*, consisting of the widely studied genus *Geobacter*, has been documented to putatively reduce insoluble Fe(III) and Mn(IV) (87) and mimicked on electrodes for anode reduction (17). This group has also been shown to uptake electron from graphite electrodes coupled with anaerobic respiration (19).

To summarize, the ISEC reactor captured a wide range of metabolically active bacteria, including (but not exclusively) well-studied groups, such as *Geobacteraceae*, shown to be capable of microbe-mineral interactions in earlier studies.

### Current production and microbial community composition in laboratory electrochemical enrichment

After 5 months of *in situ* colonization at SURF, sections of biomass-containing electrodes from ISEC reactor were transferred to laboratory electrochemical reactors. Within 8 days, cathodic currents developed at the reducing potentials (−0.19 and +0.01 V vs. SHE) and anodic currents were observed at the oxidizing potential (+0.26 and +0.53 V vs. SHE). Since the WE3 (+0.26 V vs. SHE) was lost due to an electrical short between the counter and the working electrode on the 42nd day of laboratory incubation, further analysis was performed only on WE1, WE2, and WE4 of the laboratory electrochemical enrichments. The average currents (for the last media change) corresponding to WE1, WE2, and WE4 were −43.46 ± 7.76 μA, −0.83 ± 0.06 μA, and 582.56 ± 63.14 μA, respectively (Figure 4). Separate incubation with sterile media under identical operating conditions (abiotic control) resulted in an average current of −24.976 ± 0.74 μA at WE1, −1.705 ± 0.128 μA at WE2, and 0.925 ± 0.23 μA at WE4, demonstrating that the observed cathodic or anodic currents from laboratory electrochemical enrichments are a result microbial activity on the poised electrodes. Cyclic voltammetry (CV) was performed with a scan rate of at 10 mV/s on the biofilms of colonized electrodes of WE1, WE2, and WE4 and compared to the abiotic control (Figure 4). CV of electrode-attached biofilm at WE4 revealed the classic sigmoidal shape (88) previously observed for pure EET-capable strains under turnover conditions, suggesting enrichment for biofilm containing electrode-bound, non-diffusing, redox enzymes capable of coupling oxidation of the electron donor (in this case acetate) to reduction of the electrode. Interestingly, the formal potential (as defined in (89)) of this biofilm as +0.2 V vs. SHE lies between the formal potential of *G. sulfurreducens* WT as −0.15 V vs. SHE (89) and the peak potential of outer-membrane cytochromes of *S. oneidensis* MR-1 as +0.21 V vs. SHE (90). The overall increase in potentiostatic current, the difference in the biotic cyclic voltammogram (compared to abiotic control) and cell biomass on the electrodes observed through SEM image (Figure 5), point to the electrochemical enrichment of bacterial communities capable of electron transfer to/from the electrode.

**Figure 4:**
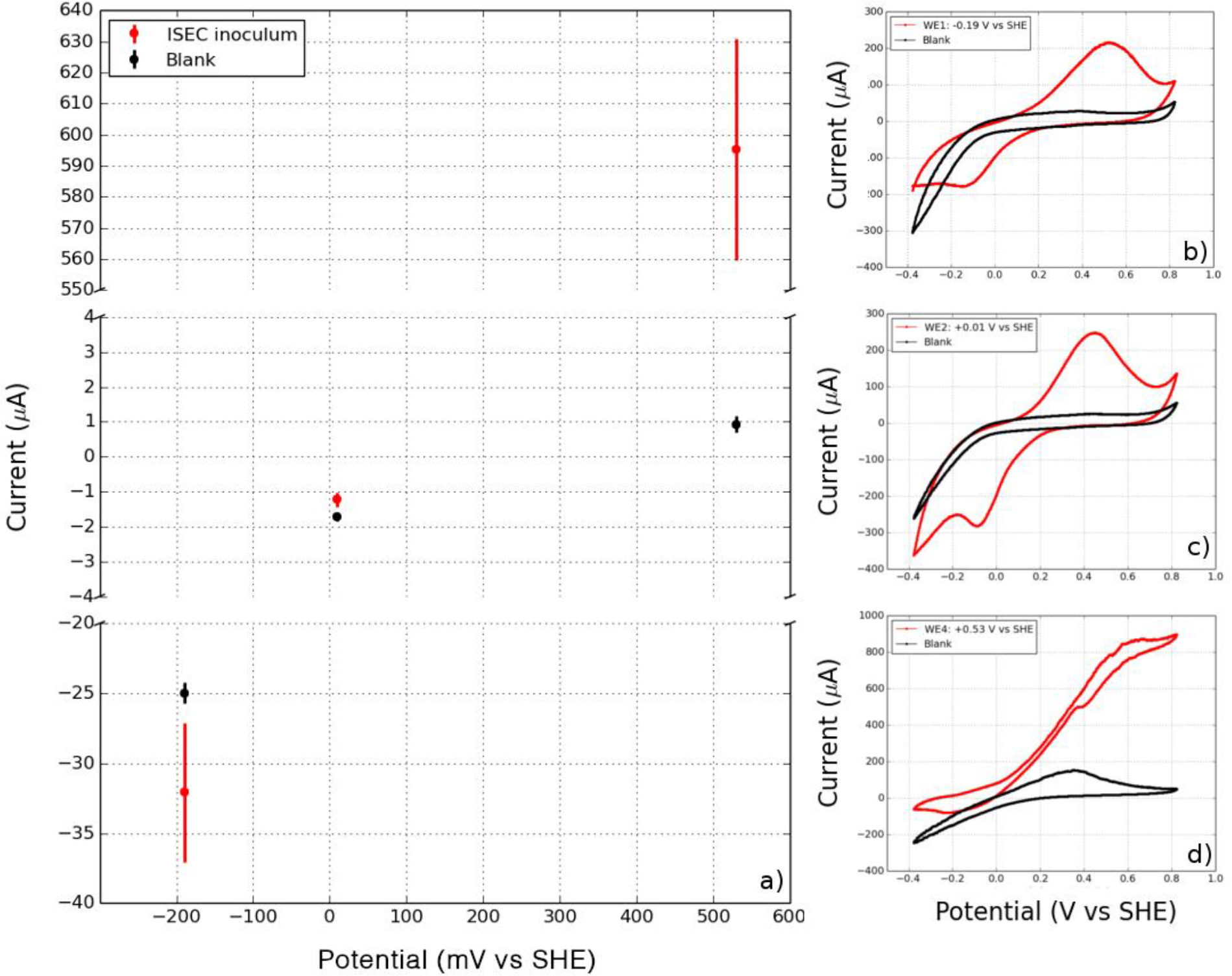
(a) Mean cathodic and anodic currents observed in laboratory enrichments at WE1 (−0.19 V vs. SHE), WE2 (+0.01 V vs. SHE) and WE4 (+0.53 V vs. SHE). Cyclic voltammetry of the working electrodes (including attached biofilms) and the corresponding abiotic controls (blank) at various potentials: b) WE1: −0.19 V vs. SHE, c) WE2: +0.010 V vs. SHE, and d) WE4: +0.53 V vs. SHE, points to redox activity mediated by the *in situ* microorganisms.

**Figure 5:**
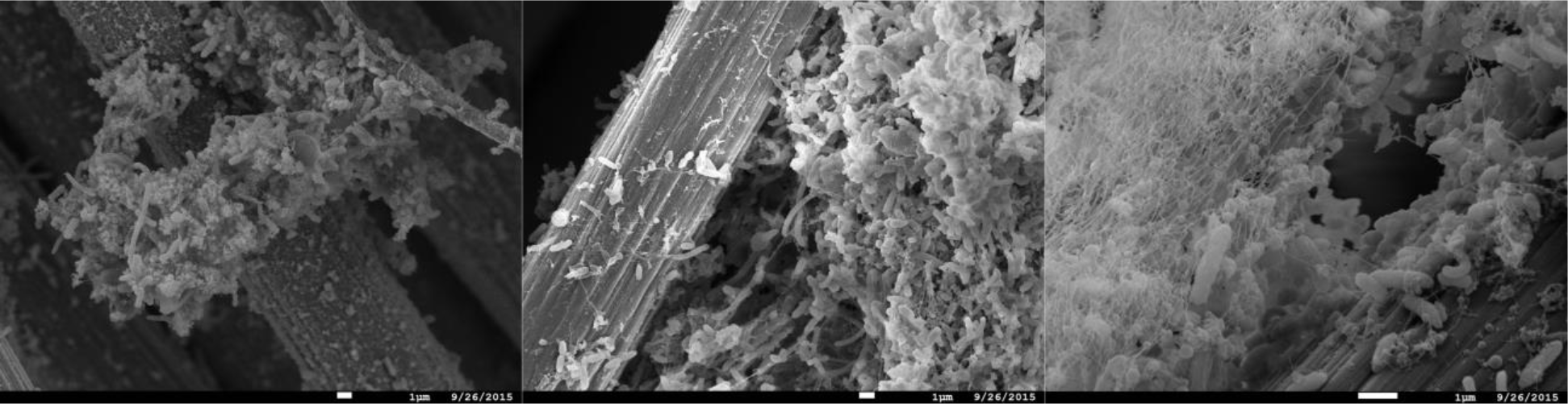
Scanning electron micrograph of the enrichment electrode-associated biomass at WE1 at −0.19 V vs. SHE (left), WE2 at +0.01 V vs. SHE (center), and WE4 at 0.53 V vs. SHE (right). (Scale = 1 μm)

The active bacterial community analysis of the laboratory electrochemical enrichments indicates growth of multiple unclassified bacterial phyla on reducing (cathodic) potentials, similar to the ISEC reactor. (Figure 6). The dominance of *Desulfuromonadaceae* and *Geobacteraceae* observed at the most oxidizing (anodic) potential of +0.53 V vs. SHE (mimicking manganese reduction) (Figure 6) explains the high anodic current and the classic catalytic sigmoidal CV signature (88). Members of these families are capable of anaerobic respiration utilizing a variety of compounds as electron acceptors, including sulfur, Mn(IV), Fe(III), and poised electrodes (91). Interestingly, *Comamonadaceae* and *Rhodocyclaceae* were present in all the poised electrodes suggesting an ability to transfer electrons to/from the poised electrodes. Many diverse bacterial families were enriched at the lower/reducing potentials (−0.19 and +0.01 V vs. SHE mimicking sulfur and iron oxidation) including the *Plactomycetaceae, Caulobacteraceae, Bradyrhizobiaceae, Hyphomicrobiaceae, Sphingomonadaceae* and *Hydrogenophilaceae*. Within *Bradyrhizobiaceae* family, some species (e.g., *Rhodopseudomonas palustris*) have been shown to oxidize Fe(II) and perform electron uptake (77) coupled with growth. The *Hydrogenophilaceae* family contains two genera, *Hydrogenophilus* and *Thiobacillus*, which are capable of gaining energy from oxidizing hydrogen and sulfur. A previous study demonstrated that the species *Thiobacillus ferrooxidans* grew by electrochemical regeneration of Fe(II) from Fe (III) at the electrode potential of 0.2 to 0.7 V vs. SHE (+0.0 to +0.5 V vs. Ag/AgCl) (92). The higher diversity of subsurface organisms at lower potential electrodes, including those that could not be unclassified at the phyla level, stress the need for further experiments to investigate extracellular electron uptake pathways.

**Figure 6:**
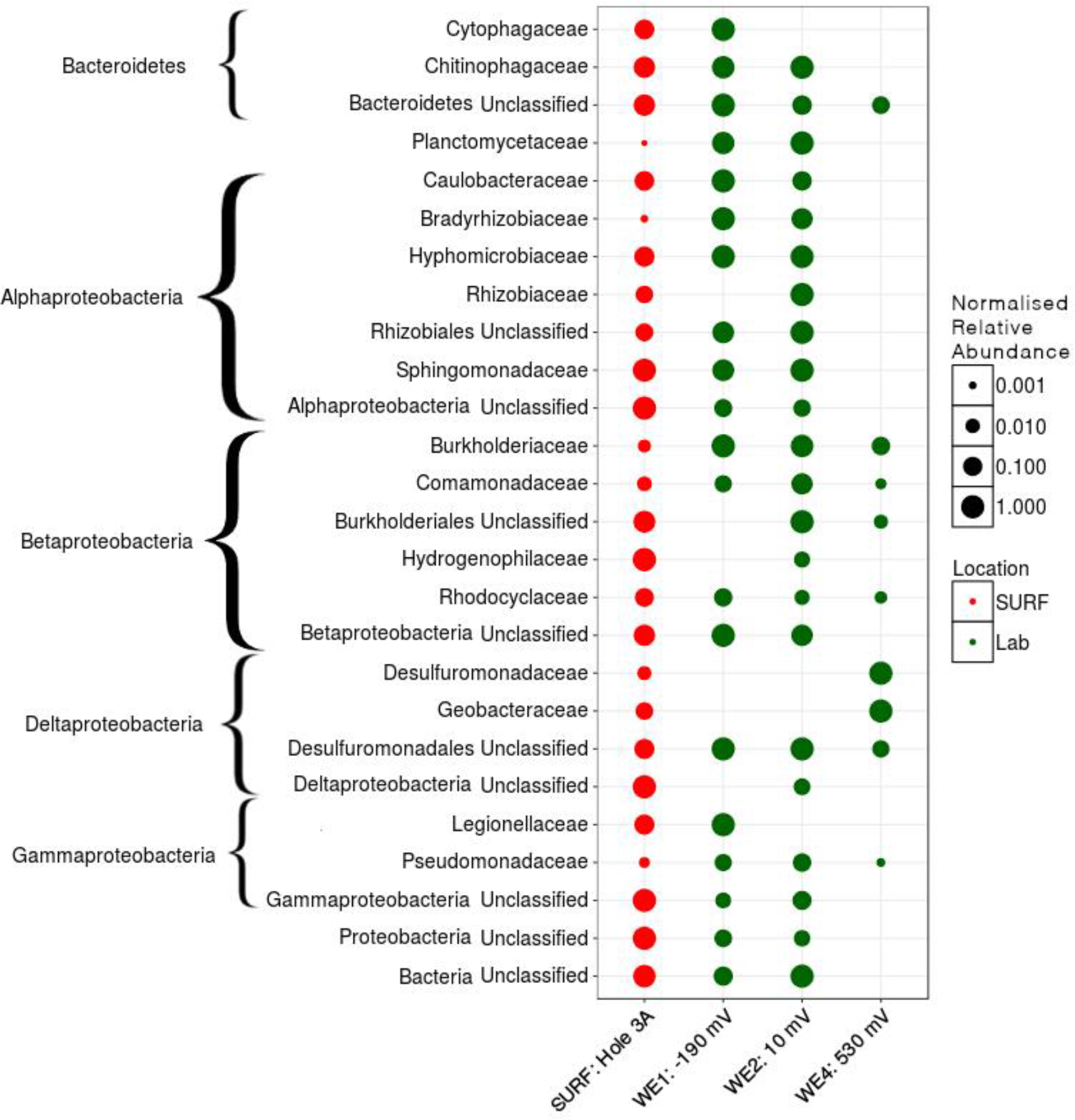
Active family-level normalized relative abundance of bacterial community present in laboratory electrochemical enrichment. The size of the symbol represents the normalized abundance of that group across various habitat types (depicted by different colors).

### Microbial community structure diversity measurements

The observed richness (number of OTUs) decreased from the source with various electrochemical enrichments (Fig 7). Nevertheless, ISEC was able to capture at least 50% of the observed OTUs at DUSEL 3A. Based on Shannon’s diversity index, there was only weak statistical evidence (Welch Two Sample T-test) of similarities in microbial community structures between DUSEL 3A and ISEC (t=3.2394, p-value=0.04235) or between DUSEL 3A and laboratory enrichment (t=3.2527, p-value=0.07365). It is interesting to note that the bacterial community attached to electrode poised at +0.01 V vs SHE (mimicking iron oxidation) has consistent higher Shannon diversity index than the communities on differently poised electrodes. OTU evenness remains constant from DUSEL 3A to ISEC and laboratory enrichment (except for the laboratory electrode poised at +0.53 V vs SHE). The consistent lower alpha diversity of biofilm enriched on the electrode poised at +0.53 V vs SHE (laboratory) is likely due to the fact that acetate was the only energy source provided. Multidimensional scaling (via PCoA analysis), evaluated using various methods, points to clustering based on location (Fig 7). Further Adonis tests performed on Bray-Curtis and Jaccard index point to variation with each enrichment step (Bray Curtis: R^2^ = 0.54918, Pr(>F) = 0.001 and Jaccard: R^2^ = 0.46127, Pr(>F) = 0.001).

**Figure 7:**
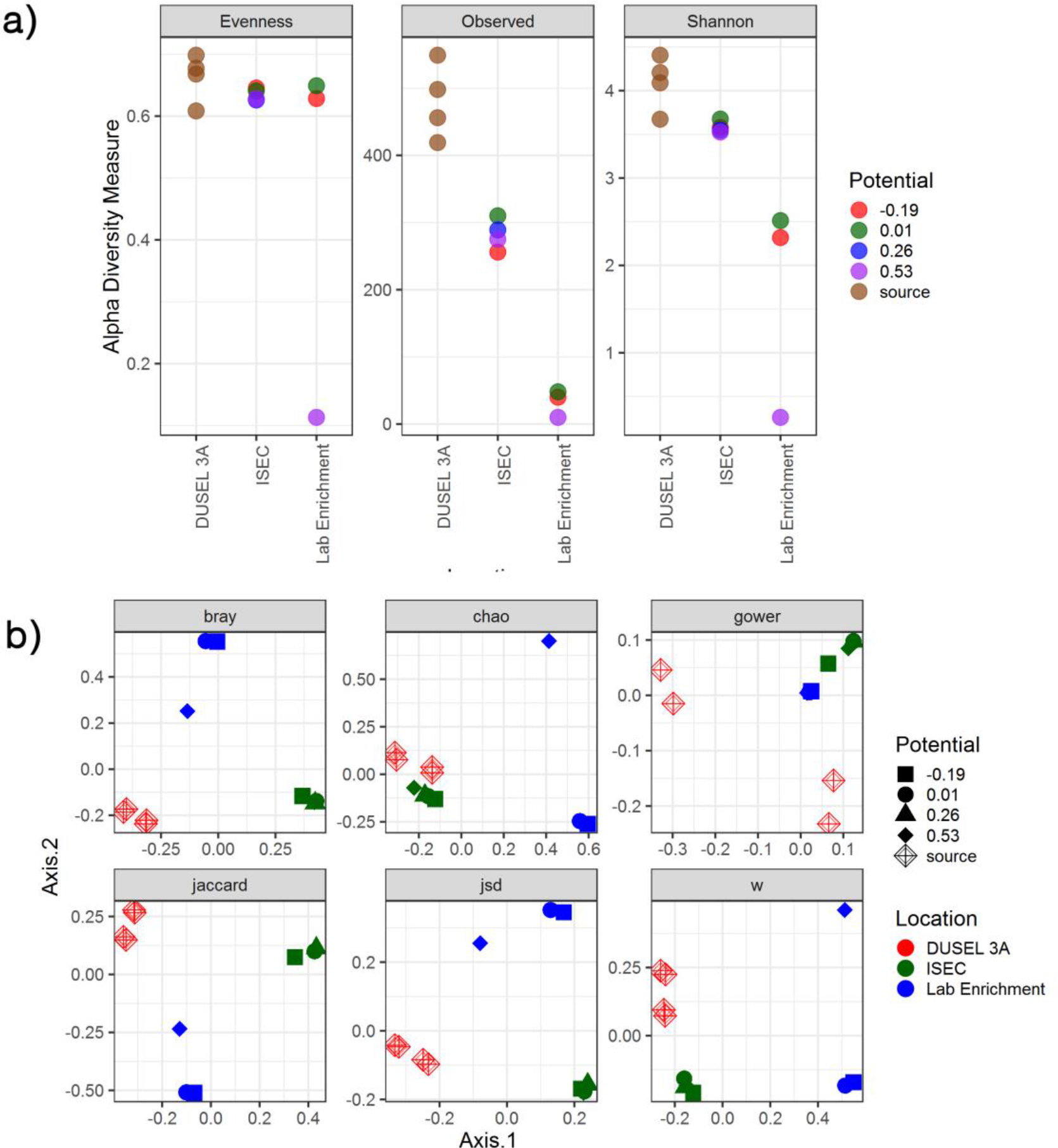
Alpha and beta diversity indices evaluated for the samples. Both the indices point to variation in the microbial community structure from ISEC to lab electrochemical enrichment. a) Estimates of evenness, richness (observed) and Shannon diversity index of the microbial community as a function of location. b) Multidimensional scaling plot evaluated using various methods points to the clustering of the microbial community based on location.

### Isolation and electrochemical characterization of isolated strains

Bacterial strains were isolated as pure cultures originating from the biomass associated with WE1 (0.19 V vs. SHE) and WE4 (+0.53 V vs. SHE). Phylogenetic analyses of the 16S rRNA gene sequences demonstrated that these strains belonged to the genera *Bacillus, Anaerospora, Cupriavidus, Azonexus, and Comamonas*. While *Bacillus, Anaerospora, Cupriavidus*, and *Azonexus* were isolated from WE4 (+0.53 V vs. SHE), two strains of *Comamonas* were isolated from both WE1 (−0.19 V vs. SHE) and WE4 (+0.53 V vs. SHE). Further electrochemical studies were performed on isolated *Comamonas* sp. (strain WE1-1D1) from WE1 (−0.19 V vs. SHE) and B*acillus* sp. (strain WE4-1A1-BC) from WE4 (+0.53 V vs. SHE). Both gram-negative *Comamonas* and gram-positive *Bacillus* have previously been reported in microbial fuel cell communities. While the genus *Comamonas* has been previously enriched on both cathodic and anodic communities of microbial fuel cells (93–95), the genus *Bacillus* has only been reported from anodic communities (96–98). Some species of these genera are capable of interacting with minerals. The *Comamonas* sp. strain IST-3 oxidizes Fe(II) for energy gain (99) and *Bacillus* can reduce Fe(III) but without conserving energy through this interaction (100).

*Comamonas* sp. (designated strain WE1-1D1) was studied in a half-cell reactor for electron uptake at working potential of −0.012 V vs. SHE. The media was constantly purged with 100% purified compressed air to supply oxygen as the electron acceptor. Upon inoculation of *Comamonas* sp. (designated strain WE1-1D1) to the reactors, the cathodic current (electron uptake from electrodes) increased from −0.51 μA to −1.67 μA (Figure 8), the planktonic cell density increased from 7 ± 1 × 10^8^ CFU/mL to 2 ± 1 × 10^9^ CFU/mL and SEM images confirmed cellular attachment with the formation of a monolayer biofilm on the WE (Figure 9). To further validate electron uptake by the strain, potassium cyanide (an electron transport chain inhibitor) was added resulting in the collapse of the cathodic current (Figure 8). Cyclic voltammetry shows a catalytic wave (reflecting cathode oxidation) with an onset potential of 50-100 mV (Figure 8). Taken together, the rise in the planktonic cell density, increase in the cathodic current, and presence of cells on the WE (Figure 9), point to electron uptake from the poised electrode by *Comamonas* sp. (strain WE1-1D1). Previously, electrochemical studies performed on the mineral oxidizing bacteria show a diversity of EET mechanisms (39–41, 44, 45). Further genomic analysis is underway to investigate the possible EET mechanism present in this strain.

**Figure 8:**
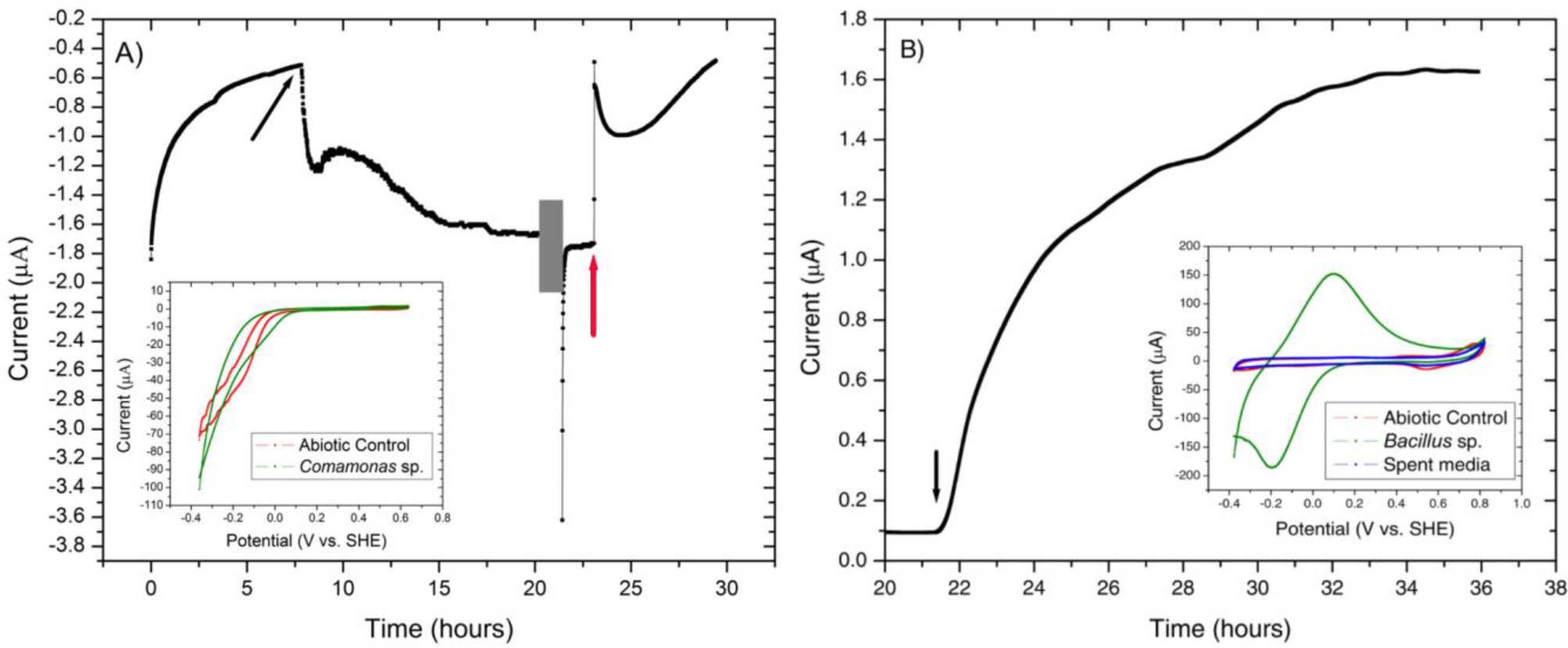
Chronoamperometry of the isolated strains (A) *Comamonas* sp. (strain WE1-1D1) and (B) *Bacillus* sp. (strain WE4-1A1-BC) grown at −0.012 V vs. SHE and +0.53 V vs. SHE respectively. A) An increase in cathodic current was observed with the addition of *Comamonas* sp. (strain WE1-1D1) (black arrow) to the electrochemical cell. At 20 hour (gray box), CV (inset) was performed on the biomass associated with the electrode and potassium cyanide was injected at 23 hours (red arrow). (B) *Bacillus* sp. (strain WE4-1A1-BC) was enriched on working electrode poised at +0.53 V vs. SHE with acetate as the electron donor and carbon source. Black arrow indicates the time point when the bioreactor was inoculated with the cells. Cyclic voltammogram of the abiotic control, attached biofilm and the spent media is shown in the inset. (SEM scale = 1 μm).

**Figure 9:**
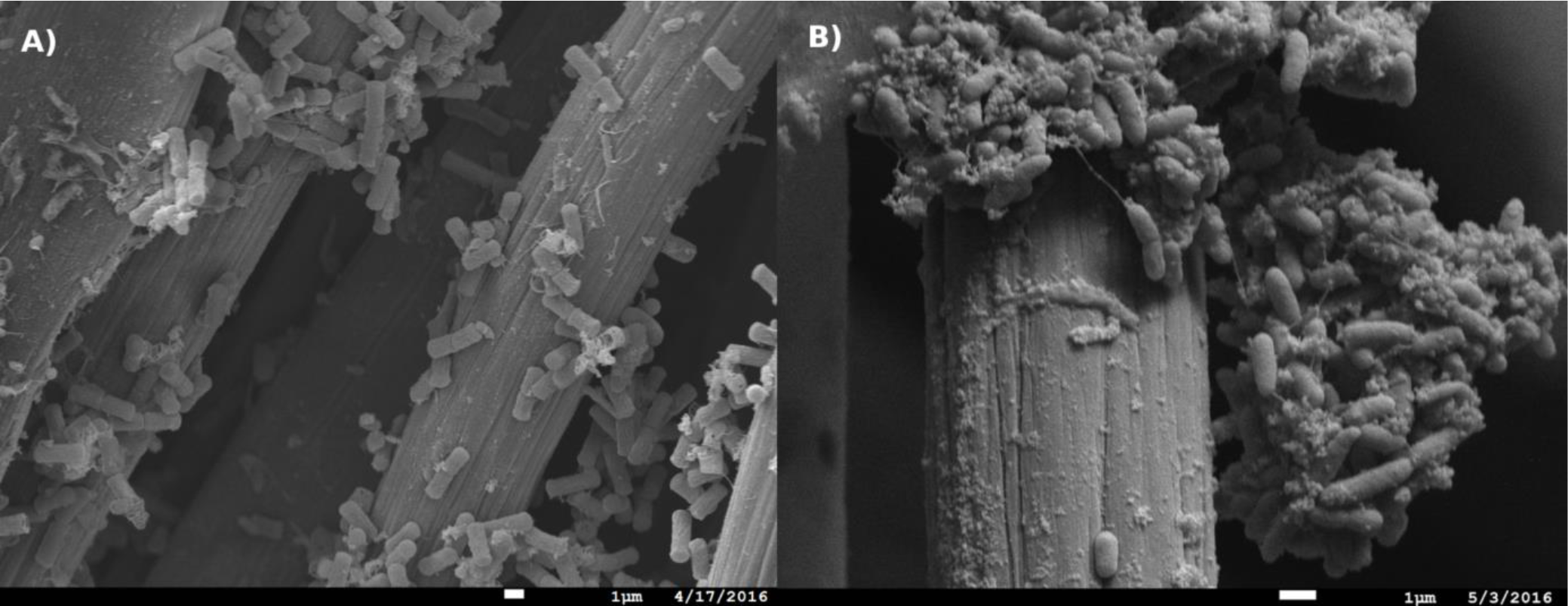
Scanning electron images of *Comamonas* sp. (strain WE1-1D1) and *Bacillus* sp. (strain WE4-1A1-BC) biomass associated with the working electrodes (Scale = 1 μm).

*Bacillus* sp. (strain WE4-1A1-BC) was studied in a half-cell reactor with working electrode poised at +0.53 V vs. SHE. Upon inoculation of *Bacillus* sp. (strain WE4-1A1-BC) to the electrochemical reactors, the anodic current rose from 0.1 μA to 1.6 μA (Figure 8) and SEM images confirmed cellular attachment with the formation of a monolayer biofilm (Figure 9). The total planktonic cell density decreased from 2 ± 1 × 10^8^ CFU/mL to 4 ± 1 × 10^6^ CFU/mL indicating possible cell lysis as free planktonic cells. Cyclic voltammetry shows a reversible peak centered at 0 V vs. SHE in the biofilm-associated working electrode. To study whether this peak arose from any soluble redox mediators released by the organism, cyclic voltammetry was performed on the spent media (planktonic cells were spun down 15× at 7000 × g). Cyclic voltammogram of the spent media also lacked the reversible peak indicating that redox components bound to the gram-positive bacteria (Figure 8) were responsible for EET. Furthermore, HPLC measurements, as described previously (101), confirmed a decrease in acetate concentration by 0.8 mM (12%) in 13 hours. Based on these measurements, the coulombic efficiency was calculated as 1.81% using the ratio of the total coulombs produced during the experiment to the theoretical amount of coulombs assuming oxidation of acetate to CO_2_. In comparison, coulombic efficiency of a well-known electrode reducer *S. oneidensis* MR-1 for electricity generation from lactate (to acetate) is ~20% (102). Taken collectively, the overall increase in anodic current, reduction in acetate concentration, the shift in biotic cyclic voltammogram relative to abiotic CV, and presence of cells on the WEs (Figure 9), suggest that *Bacillus* sp. (strain WE4-1A1-BC) is capable of electron transfer to the poised electrode. Additional studies are required to investigate the precise EET mechanism from this strain.

## CONCLUSIONS

Our results highlight the potential for poised electrodes to serve as *in situ* observatories to study and capture electrochemically active microbes in the deep terrestrial subsurface. The *in situ* electrodes deployed at the Sanford Underground Research Facility captured many families represented in the borehole fluid, while also showing potential-dependent clear shifts in the relative abundance of specific lineages. Although the ISEC reactor electrodes were poised to mimic sulfur oxidation, iron oxidation and manganese reduction, the microbial community structure pointed to sequences matching families capable of diverse metabolisms, likely reflecting interactions within and across the microbial communities colonizing separate electrochemical niches in the single cultivation vessel. Since downstream laboratory incubations were performed in separate electrochemical cells with different environmental conditions, the microbial structure of the anode-attached biofilms differed significantly from the cathode-attached biofilm. Using this workflow we were able to separately enrich and ultimately isolate cathode-oxidizing and anode-reducing microbes originating from the same environment. Further studies are needed to identify the genetic and physical mechanisms responsible for extracellular electron transfer in the isolated strains.

## ACKNOWLEDGMENTS

We acknowledge the science and support staff at SURF, including Jaret Heise, Tom F. Regan, Kathy Hart, and Oren T. Loken, for mine access, reactor deployment, and facilitating sample collection. We are also grateful to Oxana Gorbantenko and Forrest Cain, of Black Hills State University, for monitoring the ISEC reactor during the extended deployment. In addition, we acknowledge Duane Moser, Ken Nealson, Magdalena Osburn, Douglas LaRowe, Shuai Xu, Pratixaben Savalia, Lina Bird, and Annette Rowe for providing input. Scanning Electron Microscopy was performed at the University of Southern California’s Core Center of Excellence in Nanoscale Imaging. This work was supported by the NASA Astrobiology Institute (Life Underground project) under cooperative agreement NNA13AA92A. AAK and NMB were supported by the National Science Foundation DIMENSIONS grant DEB-1542527 for electrochemical characterization of the isolated strains.

